# Reconstructing the transcriptional ontogeny of maize and sorghum supports an inverse hourglass model of inflorescence development

**DOI:** 10.1101/616235

**Authors:** Samuel Leiboff, Sarah Hake

## Abstract

Assembling meaningful comparisons between species is a major limitation in studying the evolution of organismal form. To understand development in maize and sorghum, closely-related species with architecturally distinct inflorescences, we collected RNAseq profiles encompassing inflorescence body plan specification in both species. We reconstructed molecular ontogenies from 40 B73 maize tassels and 47 BT×623 sorghum panicles and separated them into transcriptional stages. To discover new markers of inflorescence development, we used random forest machine learning to determine stage by RNAseq. We used two descriptions of transcriptional conservation to identify hourglass-like developmental stages. Despite short evolutionary ancestry of 12 million years, we found maize and sorghum inflorescences are most different during their hourglass-like stages of development, following an ‘inverse-hourglass’ model of development. We discuss if agricultural selection may account for the rapid divergence signatures in these species and the observed separation of evolutionary pressure and developmental reprogramming.

**Highlights:**

- Transcript dynamics identify maize tassel and sorghum panicle developmental stages
- Random forest predicts developmental age by gene expression, providing molecular markers and an *in silico* staging application
- Maize and sorghum inflorescences are most similar when committing stem cells to a determinant fate
- Expression conservation identifies hourglass-like stage, but transcriptomes diverge, similar to ‘inverse hourglass’ observations in cross-phyla animal embryo comparisons

## Introduction

The generation of diverse organismal body plans has piqued the imagination of early naturalists and modern geneticists alike. By observing sets of intermediate developmental stages, or ontogenies, early embryologists Haeckel and von Baer not only associated the body plans of diverse species, but also placed morphological differences into the context of evolutionary relationships between taxa (Gould, 2003). Developmental genetic research has since revealed that morphogenesis generally involves a transition from a highly-proliferative stem cell identity into a determinant, mature tissue identity. By precisely regulating the duration and location of these two modes of development, multicellular eukaryotic lineages have generated complex, diverse body plans (Carroll, 2008; Minelli, 2009; Steeves and Sussex, 1972). Uncovering the molecular changes associated with the evolution of body plans underlying morphological diversity has been challenging, however, because it is difficult to determine meaningful comparisons between developmental stages of distant taxa (Roux et al., 2015), especially in understudied or morphologically ambiguous species (Anavy et al., 2014).

Our current understanding of the evolution of development has instead focused on the genetics of interfertile taxa and/or comparative genomics of species with shared morphological staging (Carroll, 2008). In systems where morphologically unique taxa are interfertile, for example, wild relatives of agricultural domesticates or allopatric species distributions, researchers have used quantitative genetics to identify mutations and even possible mechanisms underlying mutation rates that underlie morphological diversification (Hubbard et al., 2002; Jones et al., 2012; Studer et al., 2011; Xie et al., 2019). Many of these mutations provide new regulatory information for genes with important morphogenetic activity. Species which are not interfertile, but still share similar morphological staging during development can be compared with genomics techniques. For example, Lemmon et al. used morphological queues to synchronize developmental stages across species from the Solanaceae and compare transcriptomic profiles between and within genera (Lemmon et al., 2016). Comparative expression profiling approaches have similarly found that changes to the timing of expression are correlated with morphological changes (Roux et al., 2015).

Maize (*Zea mays* subsp. *mays* L.) and sorghum (*Sorghum bicolor* [L.] Monech) are two closely related cereal grains of global agricultural significance. Both members of the tribe Andropogoneae, sorghum and maize shared a common ancestor 12-16 million years ago, reflected in their extensive genomic synteny, with more than 11,000 identified maize-sorghum syntenic orthologs (Zhang et al., 2017). Despite this genomic similarity, maize and sorghum have distinct terminal inflorescence architectures, leading to differences in their agricultural use and possibly reflecting differences in their speciation/domestication histories (Lai et al., 2017; Lin et al., 2012). While much is known about the genetic underpinning of tassel morphogenesis in maize, little of that information has been applied to understanding the sorghum panicle, perhaps due to its morphological complexity (Vollbrecht et al., 2005). As interest in sorghum as a drought-tolerant biofuel and animal feed grows (Ahmad Dar et al., 2018), generating elite plant architectures will require an improved understanding of inflorescence gene function (Morris et al., 2013; Zhou et al., 2019).

Here, we present a comparative ontogeny of terminal inflorescence development in closely related grasses with morphologically unrelatable stages. By collecting individual transcriptomes from immature maize tassels and sorghum panicles throughout development, we reconstructed the transcriptional ontogeny of both species and correlated the appearance of species-specific morphological characteristics with molecularly-defined developmental stages. Examining the relative timing and sequence of known genetic master regulators from maize and their syntenic orthologs in sorghum revealed that extended tissue indeterminacy in sorghum results from the prolonged and/or heterochronic activity of multiple proliferative tissue types. We detected high transcriptional similarity between maize and sorghum during floral meristem formation, representing the termination of indeterminate, proliferative pluripotent growth. Measuring selective signatures during maize and sorghum inflorescence ontogeny detected hourglass-like mid-transition stages for each species. Despite their relatively small evolutionary distance, comparing the hourglass-like stage from each species identified their least similar transcriptional phase, providing evidence of an inverse hourglass between maize and sorghum inflorescence development.

## Results

During the formation of grass inflorescences, the pluripotent stem cells that make up the shoot apical meristem (SAM) undergo a series of proliferative tissue identity changes from indeterminate inflorescence meristems (IMs) and branch meristems (BMs), to less determinant spikelet pair meristems (SPMs) and spikelet meristems (SMs), finally terminating in completely determinant floral meristems (FMs), where all remaining stem cell initials are consumed to produce floral organs (Kellogg et al., 2013; Thompson, 2014). In maize, these stem cell identity transitions have been established by combined morphological and genetic examination of master regulatory genes (Bortiri et al., 2006; Chuck et al., 2002, 2014; Chuck and Bortiri, 2010; Eveland et al., 2014; Gallavotti et al., 2010; Thompson et al., 2009; Vollbrecht et al., 2005). Although there is extensive genomic synteny between maize and sorghum (Schnable et al., 2011; Zhang et al., 2017), inflorescence development in sorghum is sufficiently complex to be morphologically unrecognizable from maize development, a difference generally attributed to increased indeterminacy (Vollbrecht et al., 2005). We therefore used a comparative transcriptomic approach to assemble and compare complete inflorescence ontogenies for both species.

We collected individual RNAseq profiles from 40 maize tassels, inbred B73 (Figure 1A) and 47 sorghum panicles, inbred BT×623 (Figure 1B) spanning the establishment of all major architectural features. From these transcriptional profiles, we calculated complete expression trajectories for each species using a smoothing-spline pseudotime metric, developmental time units (DTUs). We then used these expression trajectories to interpolate expression values between samples and reconstruct complete molecular ontogenies (Methods; Figure 1C-F). Hierarchical clustering identified 5 transcriptional stages of maize tassel development, ZM1-ZM5 (Figure 2AB) and 4 transcriptional stages of sorghum panicle development, SB1-SB4 (Figure 2CD), sorted by DTU value.

**Figure 1.**
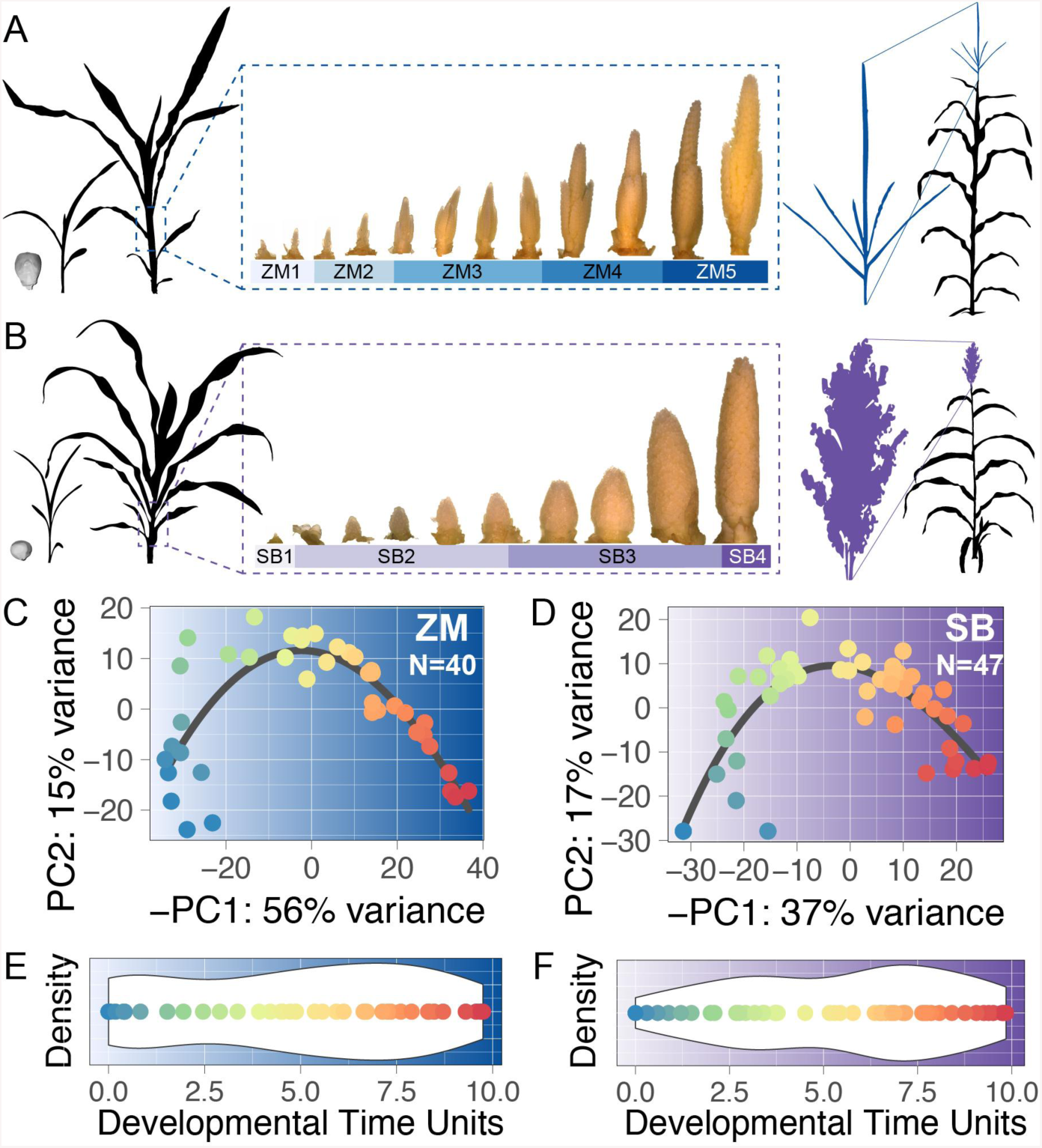
Capturing the transcriptional dynamics of inflorescence organogenesis. (A and B) Maize (A) and sorghum (B) share many features during development, but ultimately lead to divergent terminal inflorescences (right). To capture the transcriptional features of this event, we dissected, imaged, and performed RNAseq on individual inflorescence primordia (dashed box). (C and D) 5000 most-variable transcripts were used in a PCA to separate 40 maize tassel (C) and 47 sorghum panicle (D) RNAseq datasets. 2-knot smoothing splines (black line) were fitted to order and determine relative developmental progression between datasets (spectral colors). (E and F) Reconstructed molecular ontogenies for maize (E) and sorghum (F) were separated into Developmental Time Units (DTU) from 0.0 to 10.0 with relatively even representation along this developmental trajectory, white ribbon = 95% kernel smoothing density.

**Figure 2.**
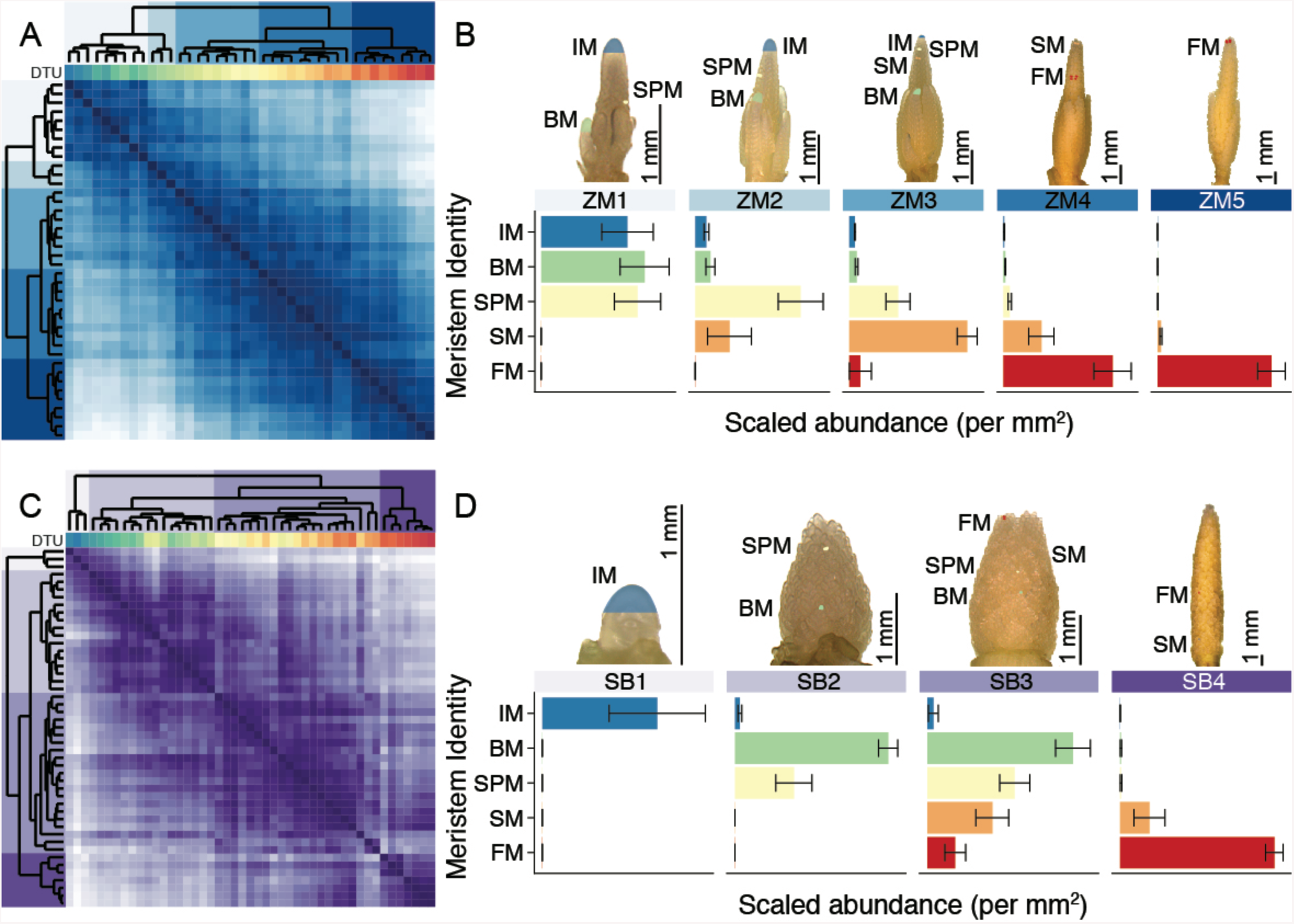
Transcriptional data identifies developmental stages correlated with changes in tissue identity. (A) Hierarchical clustering of immature maize tassel transcriptomes were sorted using DTU values (spectral colors) to produce 5 maize tassel development stages, ZM1-ZM5 (blue color bars). Pearson correlation matrix, deeper blue = higher correlation. (B) Survey of meristem types collected from ZM1-ZM5 tassel primordia before RNAseq. Average abundance of IMs (blue), BMs (green), SPMs (yellow), SMs (orange), and FMs (red) calculated per tassel area, mm^2^. ZM1 had highest proportion of indeterminate identities, IM and BM. ZM2-ZM4 were each characterized by peak abundance of SPMs (ZM2), SMs (ZM3), or FMs (ZM4) in sequence. ZM5 primordia were almost entirely FMs, although the staging of floral organs was occluded by encasing glumes. Meristem abundance ∼ stage significant by MANOVA, *F*(20, 103.77) = 20.304; p < 2.2e-16; Wilk’s Λ = 0.0050926. Univariate abundance of IM, BM, SPM, SM, and FM ∼ stage each significant by ANOVA, see Figure 2 Supplemental data 1 for test statistics. Error bars, SE. (C) Hierarchical clustering of immature sorghum panicle transcriptomes were sorted using DTU values (spectral colors) to produce 4 sorghum panicle development stages, SB1-SB4 (purple color bars). Pearson correlation matrix, deeper purple = higher correlation. (D) Sampling of meristem types collected from SB1-SB4 panicle primordia before RNAseq. Average abundance of IMs (blue), BMs (green), SPMs (yellow), SMs (orange), and FMs (red) calculated per observed panicle area, mm^2^. SB1 was the only stage significantly comprised by the IM. SB2-SB4 were each characterized by BMs and SPMs (SB2), BMs, SPMs, and SMs (SB3), or SMs and FMs (SB4), in sequence. Meristem abundance ∼ stage significant by MANOVA, *F*(15, 97.021) = 17.202; p < 2.2e-16; Wilk’s Λ = 0.027837. Univariate abundance of IM, BM, SPM, SM, and FM ∼ stage each significant by ANOVA, see Figure 2 Supplemental data 1 for test statistics. Error bars, SE.

During tissue collection, we imaged each individual inflorescence primordium, allowing the correlation of specific morphological characteristics with transcriptional phenomena (Figure 2BD). We found that our 5 maize transcriptional stages predicted the successive, acropetal (bottom-to-top) production of genetically-established meristem types, IM, BM, SPM, SM, and FM, from most indeterminate to most determinant (Figure 2B; Figure 2 Supplemental figure 1). The compound, high-order branching pattern of the sorghum inflorescence makes the panicle more spatially complex than the tassel, but stage-wise estimates of sorghum meristem type abundance matched our maize data (Figure 2D). In contrast to our maize data where the IM was observed from ZM1-ZM3, the sorghum IM was a short-lived identity found only during SB1. Subsequent meristem types appeared in a loosely basipital (top-to-bottom) sequence in sorghum (Figure 2 Supplemental figure 1).

Our calculated psudeotime metric, DTU, was tightly correlated with calendar plant age and overall primordia length, a common proxy for developmental stage (Figure 3A-D). However, we found that the relationship between DTU and sorghum panicle length could best be summarized by two piecewise linear regressions (Figure 3D), one for early panicle development (SB1-SB3) and one for late panicle development (SB3-SB4). Expression of maize meristem tissue identity genes, *faciated ear4* (*FEA4*; Pautler et al., 2015), *unbranched2* (*UB2*; Chuck et al., 2014), *ramosa1* (*RA1*; Vollbrecht et al., 2005), *branched silkless1* (*BD1*; Chuck et al., 2002), and *bearded ear1* (*BDE1/ZAG1*; Thompson et al., 2009) peaked in our dataset at DTU values that match known effects on tissue identity and published expression patterns (Figure 3E). The sorghum syntenic orthologs of these master regulator genes also predicted the appearance of different meristem types, although expression was notably shifted when comparing the two species (Figure 3F).

**Figure 3.**
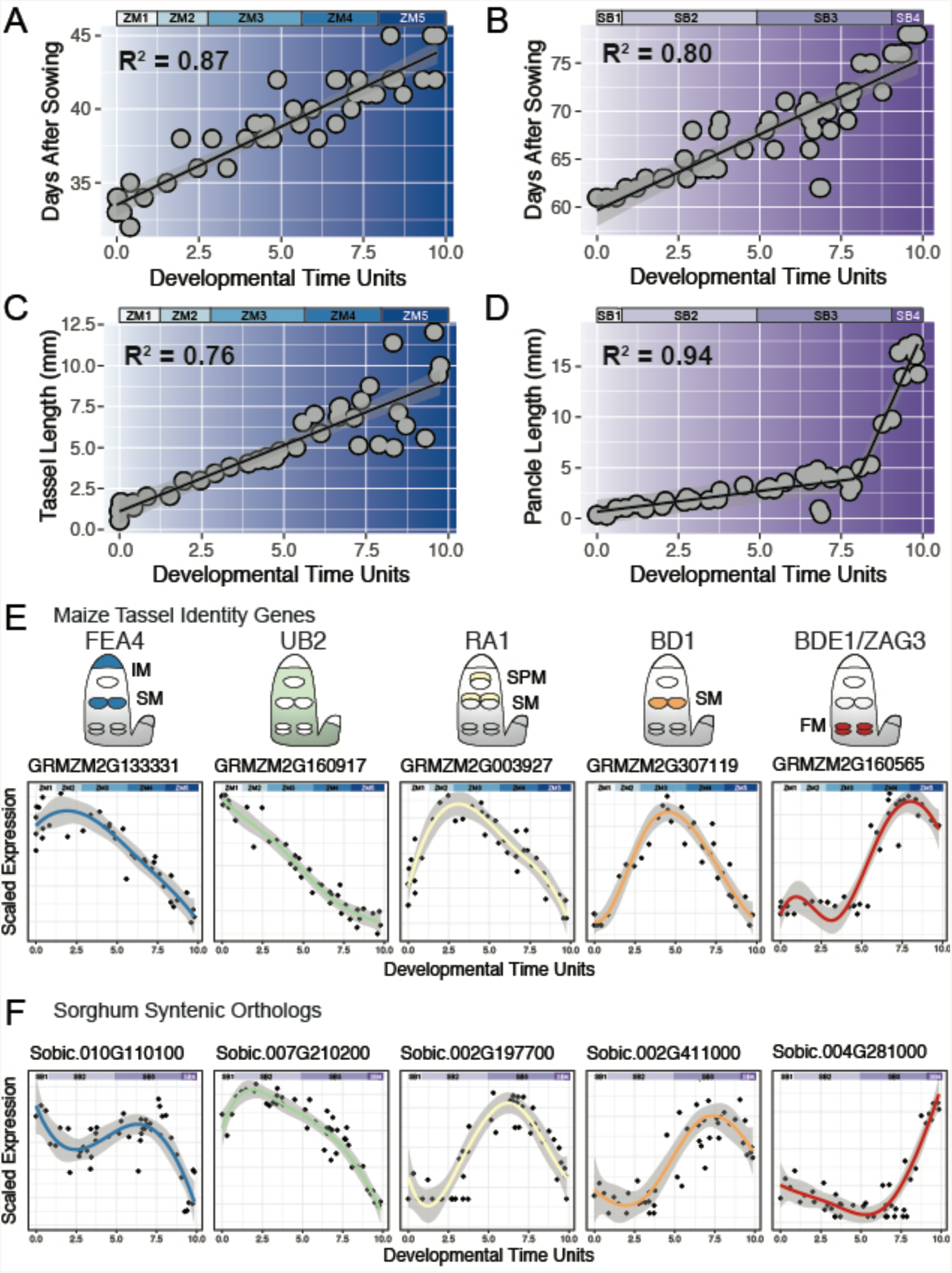
Measurements of sample age and expression of maize tassel genes and sorghum syntenic orthologs is consistent with predicted molecular ontogeny and predicted developmental stages. (A and B) Calendar age (days after sowing, DAS) was significantly correlated with transcriptionally-determined DTU in maize (A) DAS = −27.09 + 0.8262 (DTU); *F*(1, 38) = 272; p < 2.2e-16 and sorghum (B) DAS = −27.60 + 0.4841 (DTU); *F*(1,45) = 145.1; p = 1.121e-15. Regression line, black. Standard deviation residuals, grey ribbon. Adjusted R^2^ labeled in black. (C and D) Inflorescence length (mm) was significantly correlated with DTU in maize (C) length = −0.190 + 1.0022 (DTU); *F*(1, 38) = 160.7; p = 3.193e-15 and sorghum (B) length = 0.661 + 0.4089 (DTU [0,8.003)) + 7.1894 (DTU [8.003, 10]). Regression line, black. Standard deviation residuals, grey ribbon. Adjusted R^2^ labeled in black. (E) Maize tassel identity genes with functional impacts on meristem identities and their localization during tassel development (schematic of IM, SPM, SM, and FM top to bottom, BM adjacent) correlate well with peak gene expression values observed in our reconstructed molecular ontogeny. 5-knot smoothing spline, colored line. Standard deviation residuals, grey ribbon. Stages ZM1-ZM5, blue color bars. (F) Sorghum syntenic orthologs of maize tassel genes show similar trends of expression, but shifted rightward in our dataset. 5-knot smoothing spline, colored line. Standard deviation residuals, grey ribbon. Stages SB1-SB4, purple color bars.

To identify new molecular markers of inflorescence development in maize and sorghum, we employed a random forest machine learning approach that constructed decision trees based on a randomly chosen subset of gene expression values. After 2000 iterations, we calculated the informative value of each gene in predicting DTU, as a representation of developmental age. We took the top 3000 most informative genes from both maize and sorghum and clustered their expression profiles by self-organizing maps to identify stage-specific expression patterns (Figure 4A; Figure 4 Supplemental figure 1; Figure 4 Supplemental figure 2; Supplemental file 3). We then used our entrained random forest to evaluate a small number of publicly-available, developmentally-staged whole tassel primordia RNAseq datasets. Using just the raw expression values from these datasets, we were able to correctly approximate primordia length, and thus calculate organ primordia age *in silico*, from RNAseq alone for 5 of 6 available datasets (Figure 4B).

**Figure 4.**
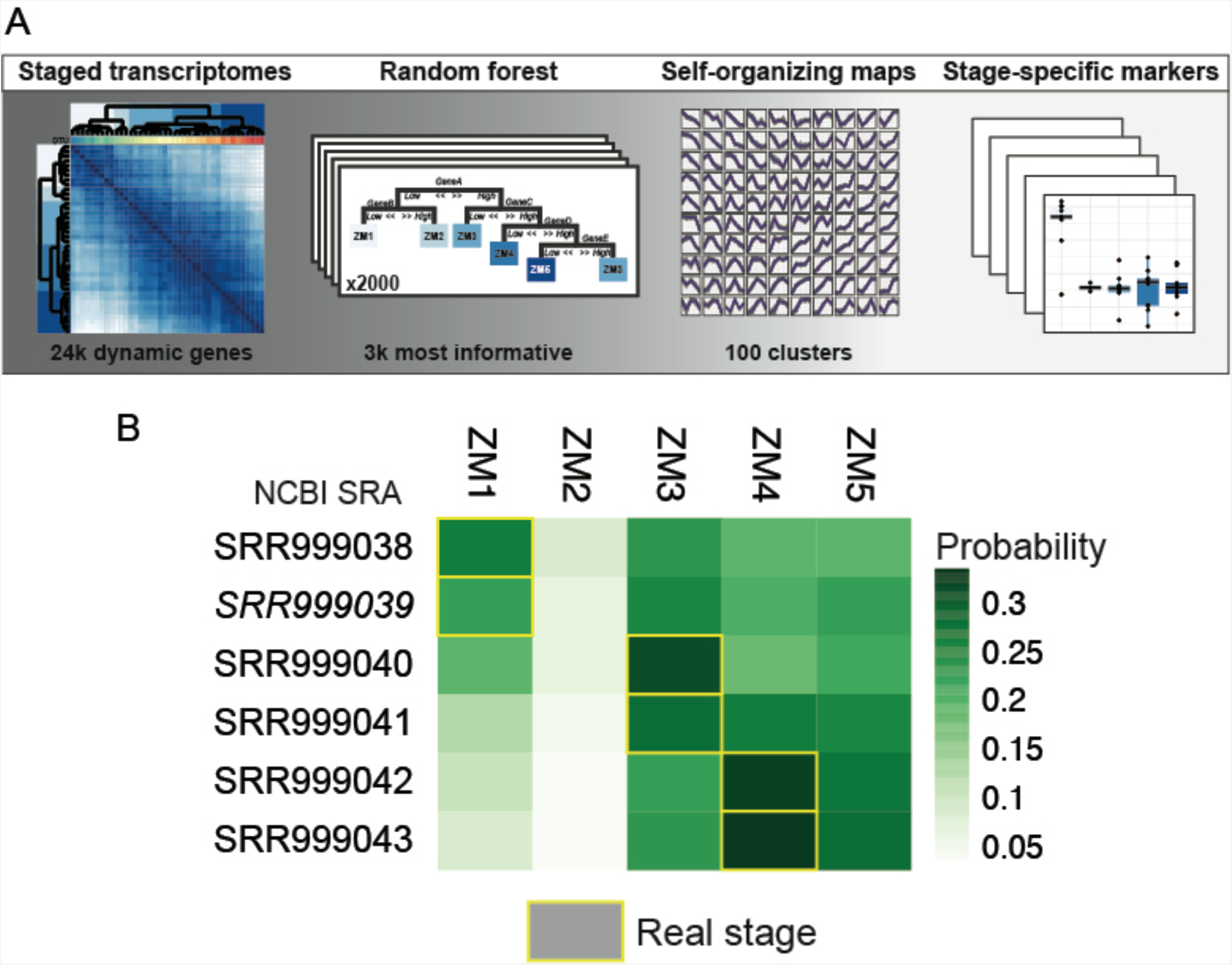
Random forest classifiers identify new inflorescence stage markers and entrain a predictive RNAseq-based model. (A) Schematic for using Random forest models to isolate stage-specific marker genes. 24k dynamically-expressed genes were used to entrain decision trees to determine sample stage by gene expression values. Using an in-bag, out-of-bag validation approach, the relative importance of each gene in determining stage could be calculated after 2000 iterations of tree-building. The top 3000 most informative genes were selected for expression profile clustering, via self-organizing maps. Genes included in clusters with stage specific expression profiles were considered as new molecular markers of maize tassel and sorghum panicle developmental stage (see supplemental tables ?? and supplemental figures). (B) Our entrained Random forest model correctly classified 5 of 6 staged, pooled maize immature maize tassel RNAseq datasets available in the NCBI short read archive. Probability of stage ZM1-ZM5, green. Actual stage inferred by reported tassel size range, yellow box.

After observing that the syntenic orthologs of maize meristem genes appeared shifted in our sorghum panicle dataset, we explored whether these shifts represent (1) differences in the relative age of sampled plants, or (2) real changes in the timing of expression, heterochrony. First we constructed phasigrams of gene expression, where genes are ordered based on their time of peak expression (Methods, after Levin et al., 2016). Comparing the absolute position of peak expression confirmed that our sorghum dataset starts and ends relatively early compared to maize development. However, we found that the overall sequence of meristem regulatory genes and their syntenic orthologs was not changed between maize and sorghum (Figure 5AB). Specific regulators associated with maize tassel development, however, did show differences in relative timing, resembling heterochrony. For example, the regulators of maize tassel branch number and complexity, *liguleless1* (*LG1*; Lewis et al., 2014) and *ramosa3* (*RA3*; Satoh-Nagasawa et al., 2006) appeared out-of-sequence, peaking later in sorghum development relative to other maize inflorescence genes. Conversely, *thick tassel dwarf1*, a negative regulator of inflorescence meristem proliferation (*TD1*; Bommert et al., 2005), peaked much earlier in sorghum development relative to maize, mirroring observations of an active IM across maize stages ZM1-ZM3 but only to the earliest SB1 stage in sorghum (Figure 2BD). We were surprised that cloned inflorescence genes displayed a limited signature of heterochrony.

**Figure 5.**
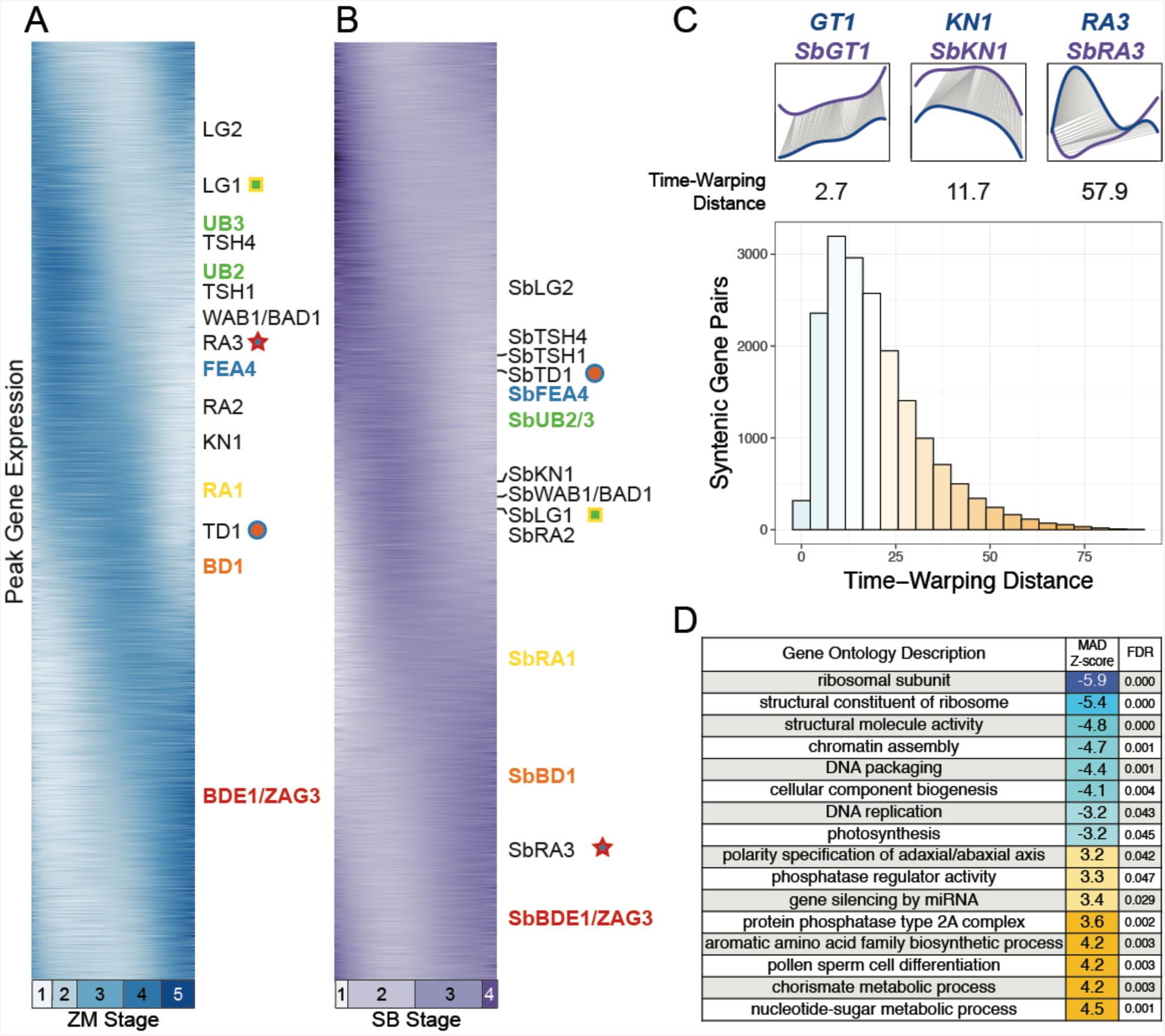
Phased gene expression schedules and Dynamic Time Warping (DTW) identify species-specific and species-shared expression profiles. (A and B) Maize (A) and sorghum (B) genes sorted by time of peak expression. Dynamically expressed genes with known tassel architecture phenotypes and their sorghum syntenic orthologs annotated. Key meristem identity genes, blue = IM associated, green = BM associated, yellow = SPM associated, orange = SM associated, red = FM associated. Syntenic ortholog pairs with notable changes in expression sequence denoted with symbols, closest ZM marker gene = symbol inner color, closest SB gene = symbol outer color. (C) DTW aligns gene expression profiles allowing for gaps, compression, and expansion, yielding a DTW distance metric for each alignment. Top, left to right, example low, near-median, and high DTW distance syntenic gene pairs. Bottom, histogram of DTW distance scores for 18k maize-sorghum syntenic gene ortholog pairs. Blue = below median DTW distance. White = median DTW distance (16.625). Orange = high DTW distance. (D) Parametric GO enrichment analysis of DTW distance median absolute distance (MAD) using maize gene annotations identified gene categories with similar (blue, negative MAD Z-score) and dissimilar (orange, positive MAD Z-score) gene expression profiles between maize and sorghum syntenic gene pairs.

To discover new genes with heterochronic expression patterns we directly compared the expression profiles of maize-sorghum gene syntenic orthologs detected in our dataset through a dynamic time warping (DTW) profile-alignment metric (Giorgino, 2009). DTW compares expression profiles, allowing gaps, compression, and expansion of one gene expression profile in order to fit another (Figure 5C). Genes with low DTW distances have similar expression profiles, even if they are expressed at different absolute times. Genes with high DTW distances have dissimilar expression profiles and cannot be synchronized by simple translation. Syntenic maize-sorghum orthologs varied from low to high DTW, with a bias towards low DTW distances (Figure 5C). We used median absolute deviation normalized DTW scores in a parametric gene enrichment test to search for enriched GO terms within similar and dissimilar expression profiles (Supplemental file 4). Genes annotated with GO terms related to DNA replication, regulation of photosynthesis and other core processes were amongst the most similar between maize and sorghum inflorescence development (Figure 5D), suggesting that general features of growth are regulated similarly in maize and sorghum inflorescence development. On the other hand, GO terms related to adaxial-abaxial specification, secondary metabolism, and PP2A complex were enriched in the most dissimilar maize-sorghum gene expression comparisons, suggesting that floral organ programs are amongst the most heterochronic expression patterns between species.

To assess global similarities in gene expression, we limited our gene expression dataset to syntenic maize-sorghum orthologs, divided each reconstructed developmental expression profile into 1000 time points, and calculated Pearson correlation between maize and sorghum inflorescence development in all pairwise combinations (Figure 6A). By comparing the full trajectory of gene expression during maize tassel and sorghum panicle development, we detected continuous transcriptional similarity between maize and sorghum. We found low transcriptional similarity during the appearance of spikelet pair meristems in maize and peak branch meristem abundance in sorghum (ZM2, SB2) and high similarity during floral meristem accumulation in both species (ZM4, SB3; Figure 6A). Maize and sorghum inflorescence development are least similar when comparing ZM4, marked by high floral meristem abundance to SB2, marked by peak branch meristem abundance (Figure 6A). We further used this similarity to assemble a linear relationship between maize tassel and sorghum panicle developmental stages (Figure 6B).

**Figure 6.**
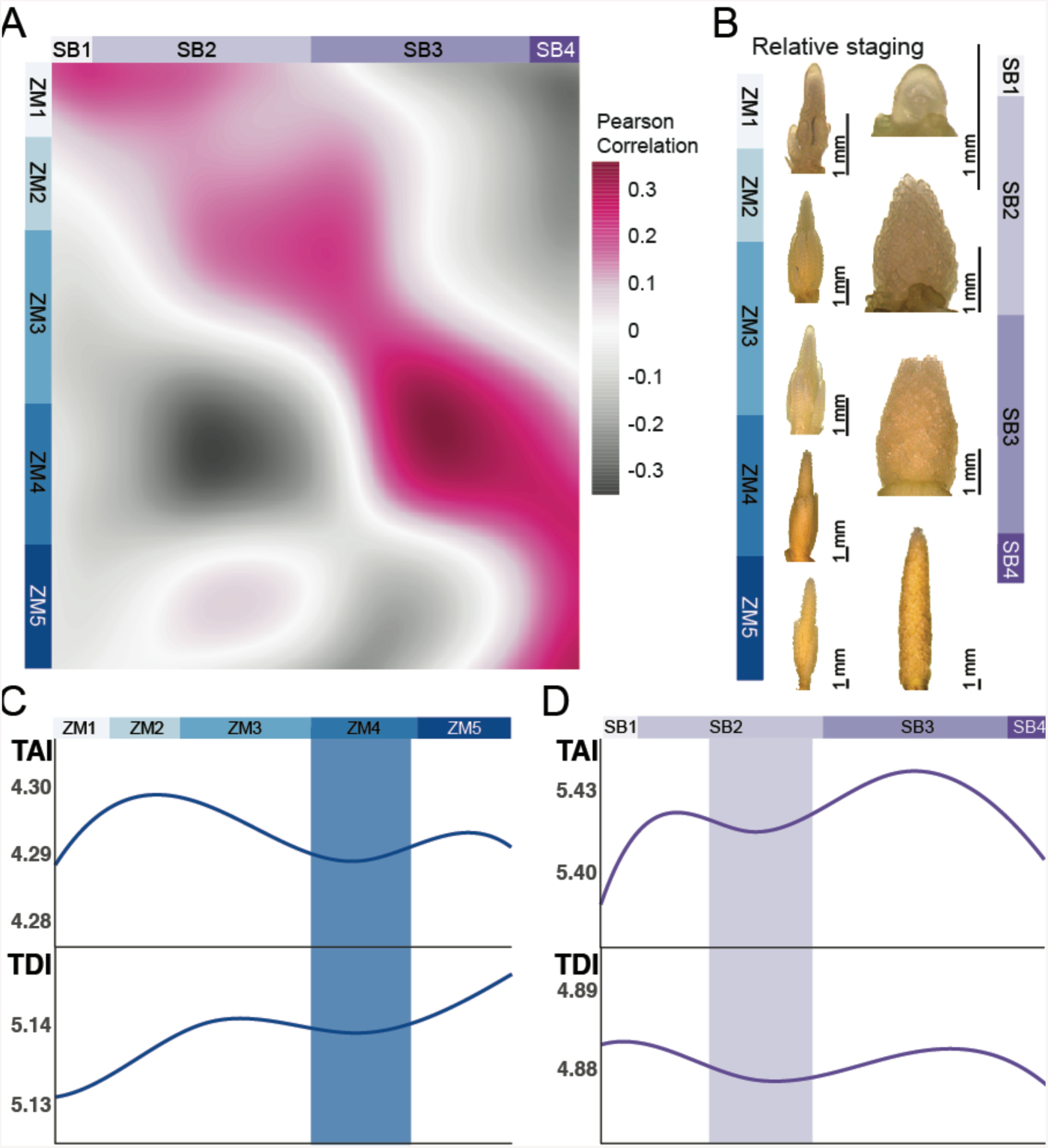
Synchronizing maize and sorghum inflorescence development finds dissimilar hourglass-like developmental stages, as expected in an inverse-hourglass model of development. (A) Correlation between maize and sorghum inflorescence molecular ontogenies. Peak correlation during ZM4 and SB3, both times with high FM activity, and minimal correlation during ZM4 and SB2. Maize stages, ZM1-ZM5 vertical blue color bars. Sorghum stages, SB1-SB4 horizontal purple color bars. Pearson correlation, positive = pink, no correlation = white, negative = grey. (B) Tracking peak correlation between maize and sorghum expression dynamics determined relative staging. Maize stages, ZM1-ZM5 blue color bars. Sorghum stages, SB1-SB4 purple color bars. (C and D) Signatures of anciently conserved transcriptional, developmental hourglass-like stages by phylostratigraphy (TAI) and codon divergence stratigraphy (TDI); maize ZM4, blue vertical band; sorghum late SB2, purple vertical band. Comparing these stages by overall transcriptome similarity (A) suggests an inverse-hourglass relationship between maize tassel and sorghum panicle development. (C) TAI significant by flat line test, p = 3.22e-05, TDI significant by flat line test, p = 2.68e-07. (D) TAI significant by flat line test, p = 1.61e-45, TDI significant by flat line test, p = 5.69e-03.

Developmental expression profiling has revealed that animal, fungal, and plant transcriptomes exhibit signatures of a ‘developmental hourglass’ where the ‘hourglass-like’ stage is enriched for the expression of anciently-conserved genes (Cheng et al., 2015; Drost et al., 2015; Quint et al., 2012). This hourglass-like stage coincides with the establishment of an organism’s body plan and is morphologically conserved within closely-related species. Seeking to understand the source of maize and sorghum morphological differences from our transcriptomic data, we used an evolutionary transcriptomics approach to explore whether there is an analogous ‘developmental hourglass’ during maize and sorghum inflorescence development.

We employed two approaches to understand transcriptional conservation. We combined phylostratigraphy, where the ancestry of maize peptides was inferred by protein BLASTs against a representative set of 216 plant, animal, and bacterial genomes with our developmental transcriptomes to produce a transcriptional age index, TAI (Arendsee et al., 2019; Drost et al., 2018; Quint et al., 2012). In parallel, we determined the conservation of maize or sorghum codons by calculating Ka/Ks in reciprocal best-BLAST-hits against the *Setaria italica* genome in a codon divergence stratigraphy approach to determine a transcriptional divergence index or TDI (Drost et al., 2018; Quint et al., 2012). Across maize tassel development, TAI and TDI fluctuated significantly, with an increased contribution of anciently-conserved genes during high floral meristem abundance detected by both phylostratigraphy and codon divergence stratigraphy (ZM4; Figure 6C). The relative increase in ancient gene activity was driven by the expression of genes shared amongst all green plants (Streptophyta, Viridiplantae) as well as gene modules shared amongst most monocots (Petrosalviidae, commelinids, Poales; Figure 6 Supplemental figure 1). However, the absolute expression value of maize-specific and Andropogoneae-specific genes was greater than all other strata at all maize tassel transcriptional stages (ZM1-5; Figure 6 Supplemental figure 1). Across sorghum panicle development, we detected a significant increase in ancient-conserved genes shared amongst monocots (Liliopsida, Petrosaviidae, commelinids) with a less prominent contribution from genes shared amongst all green plants (Viridiplantae, Streptophyta, Embryophyta; Figure 6 Supplemental figure 1) during the proliferative branching stage (SB2). As seen in maize, absolute expression values were dominated by sorghum-specific and Andropogoneae-specific transcripts during all sorghum panicle stages (SB1-4; Figure 6 Supplemental figure 1).

Although we detected hourglass-like signatures of purifying selection of transcriptional programs in maize tassel stage ZM4 and sorghum panicle stage SB2, morphological comparisons suggest that these stages are not analogous, with ZM4 representing the specification of determinant floral meristem identity and SB2 comprised of highly indeterminate compounding branch meristems (Figure 2CD). And while transcriptional stages, ZM4 and SB3 exhibited high Pearson correlation, our putative hourglass-like stages, ZM4 and SB2 display low Pearson correlation, matching expectations for a developmental ‘inverse hourglass’ normally detected in distant species with dissimilar body plans (Lemmon et al., 2016; Levin et al., 2016; Yanai, 2018).

## Discussion

As a way of understanding the similarity of the middle stages of vertebrate embryogenesis, Duboule’s concept of the phylotypic egg timer, also known as the ‘developmental hourglass’, pointed to mechanistic constraints on development as a source of shared morphology (Duboule, 1994). Duboule proposed that the linked, tightly regulated clusters of colinearly-expressed HOX genes force vertebrate embryos into a similar body plan. Strong selection for this developmental mechanism and the body plan it produces thus underlie the similarity in limb-bud stage of development in both zebrafish and mice, vertebrates that shared an ancestor more than 400 MYA. Exploring the embryonic mid-transition of closely-related animal species has found genome-wide molecular evidence for a conserved transcriptional program within phyla, where the activity of evolutionarily-conserved genes establishes characteristic body plan features shared within the phylum and is maintained by strong purifying selection at the ‘hourglass-like’ stage (Anavy et al., 2014; Drost et al., 2015; Kalinka et al., 2010; Levin et al., 2012). Simulations of gene network evolution suggest that changes to developmental pathways can quickly define an hourglass-like stage through the loss of expression of formerly-interacting genes (Akhshabi et al., 2014). Duboule suggested that insects, which share a common ancestor with vertebrates at least 530 MYA, do not experience this same developmental constraint because insect lineages have multiple HOX clusters that are not globally, colinearly expressed and thus insect and vertebrate embryogenesis is morphologically and mechanistically distinct (Duboule, 1994).

While a transitional stage between stem cell proliferation and tissue patterning is detectable across diverse taxa, broad comparisons between metazoan phyla (Levin et al., 2016), plant species within the same family (Lemmon et al., 2016), or between animals, fungi, and plants (Cheng et al., 2015; Drost et al., 2015; Quint et al., 2012), reveal that although hourglass-like mid-transition patterns of conservation can be detected, evolutionarily-distant taxa have dissimilar mechanisms regulating the mid-transition. Wide transcriptomic and morphometric comparisons of anciently diverged taxa have thus lead to an ‘inverse-hourglass’ model of cross-phyla development, where the greatest dissimilarity between species is detected by comparing their unique ‘phylotypic’ mid-transitions, with very few developmental mechanisms conserved at the mid-transition across phyla (Levin et al., 2016).

Maize and sorghum, as members of the tribe Andropogoneae, are estimated to have shared a common ancestor as early as 12 MYA. The maize tassel and sorghum panicle share characteristic morphological features, including a branched inflorescence terminating in short paired spikelet branches bearing paired florets. These features are shared amongst the Andropogoneae and other grasses (Kellogg et al., 2013). In this study we detected clear continuous linear transcriptomic correlations between maize tassel and sorghum panicle developmental stages, suggesting that the bulk of their developmental activities are shared. Despite these general similarities in body plan, short evolutionary history, and correlated expression patterns, phylostratigraphy and codon divergence stratigraphy suggest that a relatively short history of selective pressures have allowed the hourglass-like stage to reposition in either or both of these species (Figure 7A). Without knowing the expression dynamics of their common ancestor, or a comprehensive panel of other grasses, we cannot determine whether maize-like selective pressure on FM specification or sorghum-like selection on BM indeterminacy marked their last shared ancestor. Indeed, the last ancestor of maize and sorghum may have exhibited a completely different hourglass-like stage of inflorescence development. In any case, our findings suggest that selective pressures have acted differently on maize and sorghum, leading to a change in the hourglass-like signature of selection independent of developmental diversification (Figure 7B).

**Figure 7.**
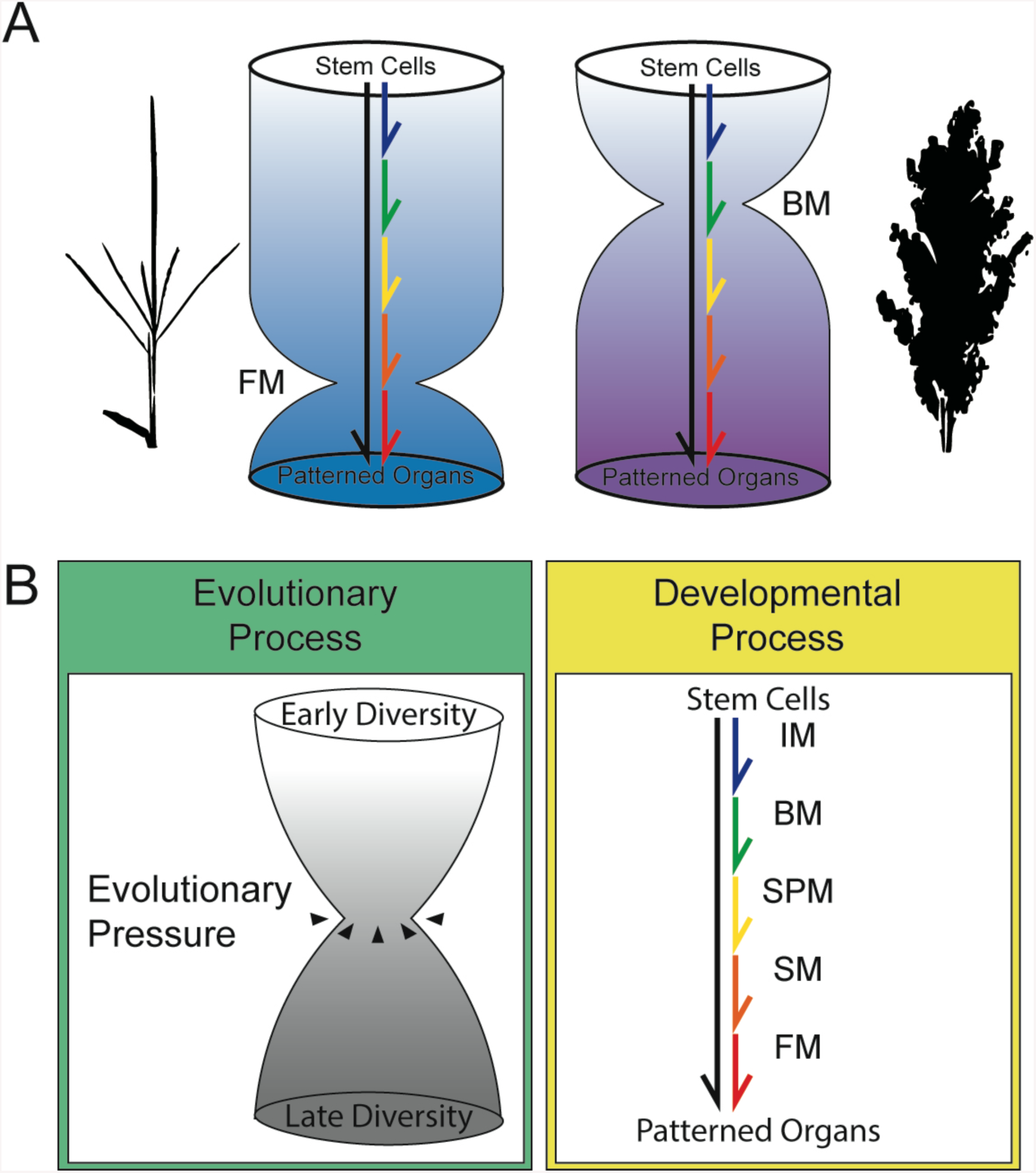
Differences in hourglass-like stages in maize and sorghum inflorescences suggest that evolutionary and overall developmental processes can be uncoupled. (A) Hourglass-like stages occur during maize tassel development when determinant floral meristems, FMs are most abundant (left), but also during sorghum panicle development when indeterminant branch meristems, BMs are most abundant (right). Overall signatures of inflorescence development are similar, arrows. (B) Observed separation in hourglass-like stage definition by evolution, where evolutionary pressure restricts gene activity by purifying selection (left), yet organogenic pathways are independently refined (right).

We wondered if domestication, which acts as a strong allelic bottleneck that rapidly changes morphology, could be the driving force behind the observed differences in the maize and sorghum inflorescence hourglass-like stage. We searched for enrichment of genes associated with maize domestication (Hufford et al., 2012) amongst our calculated phylostrata and codon divergence strata, but did not find a significant enrichment in any stratum (Supplemental file 5), suggesting that maize domestication does not play a significant role in defining the maize hourglass-like stage. Additional investigations into the genome-wide signatures of domestication in sorghum would allow us to further determine whether agricultural selection is sufficiently powerful to reprogram the developmental hourglass without disturbing global molecular ontogeny.

## Conclusion

By collecting individual transcriptional profiles that span inflorescence maturation, we reconstructed a complete molecular ontogeny of maize tassel and sorghum panicle development. We used our molecular ontogeny to identify 5 maize tassel and 4 sorghum panicle developmental stages. These stages correlated with quantitative morphological signatures of tissue identity, although sorghum and maize inflorescence displayed different spatial distributions of tissues.

As seen in models trained to identify novel biomarkers (Scheubert et al., 2011), we were able to use machine learning (ML) algorithms to identify important transcriptomic features. That the entrained model is able to perform *in silico* sample staging with other RNAseq data suggests that carefully prepared ML models might one day allow naive users to characterize samples with the same precision.

Although our transcriptional data identified widespread molecular similarities in maize tassel and sorghum panicle development, a known regulator of determinacy, *RA3*, and stem cell homeostasis, *TD1*, are not expressed in a shared pattern during inflorescence development in both species. Our data support earlier reports that increased indeterminacy via heterochrony is correlated with the high-order branching of the sorghum panicle, compared to the maize tassel (Vollbrecht et al., 2005), but that this process is mediated by few known master regulatory genes. The genome-wide characterization of heterochrony by dynamic time warping (DTW) promises to reveal new genes underlying morphological differences in maize and sorghum inflorescences.

By identifying hourglass-like stages in maize tassel and sorghum panicle development we show that developmental hourglass patterns of embryonic similarity may be applicable to post-embryonic phases of plant development, along with development in the seed (Drost et al., 2016; Quint et al., 2012), supporting a life-long iterative, modular rhythm to plant development (Kaplan and Cooke, 1997). That we detected changes in hourglass-like selective signatures between maize and sorghum which were not in agreement with transcriptome-wide similarities suggests that relatively brief evolutionary pressures may influence hourglass-like signatures while leaving broad developmental mechanisms intact. These data suggest that forces measured across phylogeny need not be recapitulated in ontogeny.

## Acknowledgements

We thank Z. Lemmon and C. Marklez for early discussions that lead to this work. SL is supported by NSF IOS 1612268 NPGI Postdoctoral Fellowship. This work used the Vincent J. Coates Genomics Sequencing Laboratory at UC Berkeley, supported by NIH S10 OD018174 Instrumentation Grant.

## Methods

### Tissue collection, imaging, and RNA profiling

*Zea mays* subsp. *mays* inbred B73 (PI 550473) and *Sorghum bicolor* inbred BT×623 (PI 564163) were grown in three staggered plantings in greenhouse conditions. Individual kernels were sown in 36-well starter trays every week for three weeks. Germinated maize seedlings were transplanted to 2-gallon pots, 4-to-a-pot after one week. Germinated sorghum seedlings were transplanted to 2-gallon pots, 4-to-a-pot after two weeks. Plants from one pot were harvested from 2-4 PM every day from all three plantings from 30 DAS to 80 DAS as needed.

Plants were dissected with the aid of a Leica MZ 16 F stereoscope and imaged using a Teledyne QImaging MicroPublisher 6 CCD camera at 1× and 5× magnification. Leaves were removed with a scalpel to reveal the developing primordium. During imaging, plants were ranked 1-5 stars based on the integrity of the primordium, areas of damage, dust, etc. Only samples with 4-5 stars were used to produce RNA and cDNA sequencing libraries. Immediately after imaging, each primordium was sealed in a 1.5 ml eppendorf tube containing 4 chromium beads and flash frozen in liquid nitrogen while other plants were harvested. Dissections that required more than 30 minutes to complete were discarded to reduce the chance of tissue damage related transcriptional changes.

After harvest, tubes were removed from liquid nitrogen and homogenized using a Retsch MM 301 tissue homogenizer (freq = 30 Hz, duration = 30 sec), then returned to liquid nitrogen. RNA was extracted using TRIzol (Invitrogen), precipitated using 0.5M NaCl + 2-propyl alcohol, washed with 70% ethanol, and resuspended in nuclease-free water. RNA was quantified using a QuBit BR-RNA kit and quality was evaluated on a 1% agarose gel in 1× TAE buffer.

Approximately 1 ug of RNA was used as input to the NEB Ultra II RNA sequencing library kit for illumina (New England Biolabs, NEB catalogue number: E7775L, E7335S, E7500S, E7710S, E7730S). Multiplexed library synthesis was carried out according to manufacturer’s specifications, making use of optional poly-dT bead mRNA selection (NEB E7490L) and 7-cycle PCR amplification as specified. Library quality and quantity was verified by Agilent DNA BioAnalyzer and qPCR by the QB3 Vincent J. Coates Genomics Sequencing Laboratory.

### RNA sequencing, alignment, and normalization

Maize and sorghum libraries were added in equimolar mixes and sequenced separately, each using one lane of an Illumina HiSeq4000 with 100 bp single-end sequencing chemistry. Sequence quality was evaluated using FastQC and MultiQC. Illumina adapter sequences were trimmed using Trimmomatic. Reads were aligned to the maize B73 AGPv3.30 or sorghum BT×623 v3.0.1 genome using HiSAT2. Aligned reads were counted using a union-exon approach with HTseq-Counts to the B73 AGPv3.30 gene set or BT×623 v3.1.1 gene set. Raw counts were normalized using variance-stabilizing normalization with DESeq2. Genes with less than 5 reads per million or detected in less than 39 sequencing libraries were not considered in subsequent analysis.

### Sample psuedotime indexing and stage determination

We used row variance to identify the top 1500 most variable genes and separate samples based on a principal component analysis (PCA) for both maize and sorghum datasets. A 3-knot b-spline was fit to component 1 and component 2. Each sample was assigned a location on the b-spline by minimizing the Euclidean distance between the spline and the real expression dataset. The rank and distance along the b-spline was used to calculate a Developmental Time Units (DTU) value from 0.0 to 10.0.

The complete gene expression matrix was used for hierarchical clustering by average linkage and produce a dendrogram of between-sample relationships with R dendextend (Galili, 2015). DTU was used to sort the branches. Stages were determined by clustering distance.

### Image analysis

Tiled 1× and 5× images from harvested tissues used for RNAseq were analyzed using ImageJ (Schneider et al., 2012). Meristem tissue identity was determined by appearance, counted, and quantified using the count objects tool. Binary thresholding was used to determine the area of the total inflorescence silhouette and normalize meristem abundance by inflorescence size. Late stage sorghum samples were too large to survey completely, so we quantified 5 randomly positioned image subsamples and used the subsample silhouette to normalize abundance by size.

### Random forest modeling, prediction, and clustering

The randomForest package for R was used to entrain a random forest model to predict inflorescence stage with an unfiltered, variance-stabilized gene expression matrix (Liaw and Wiener, 2001). The optimal number of trees and number of variables at each split point were determined empirically by minimizing out-of-bag error rates, maize: ntree=2000, mtry=106; sorghum: ntree=2000, mtry=4.

B73 tassel datasets were accessed from the NCBI SRA (BioProject PRJNA219741; accession SRR999038, SRR999039, SRR999040, SRR999041, SRR999042, SRR999043). Tassel stage was predicted using the entrained random forest and aggregated stage assignment probabilities were reported.

The decrease in accuracy for each gene feature during random forest model entrainment was used to identify the top 2500 most influential genes. These most influential genes were clustered using self-organizing maps with a 10 × 10 hexagonal grid and 50,000 iterative steps (R package kohonen; Wehrens and Buydens, 2007; Wehrens and Kruisselbrink, 2018).

### Expression comparison of syntenic orthologs

The variance stabilized gene expression matrix was used to fit a 5-knot b-spline for each maize and sorghum gene. Each gene’s fitted curve was interpolated into 1,000 points along its expression trajectory to allow for smooth, continuous comparisons. To produce phasigrams (Levin et al., 2016), we performed PCA on z-scaled expression values for each gene. When plotted, component 1 and component 2 formed a circle. We used the atan2 function to order genes based on their time of peak expression. Maize genes and their sorghum syntenic orthologs (Zhang et al., 2017) were identified as annotations on a vertical heatmap based on atan2 ordering.

Dynamic time warping (DTW) was performed on maize-sorghum syntenic ortholog pairs using z-scaled expression values and the R package dtw (Giorgino, 2009). Median absolute deviation was used for parametric gene enrichment tests (Tian et al., 2017) of species-shared and species-specific expression patterns.

### Evolutionary expression analysis

We calculated maize and sorghum peptide phylostrata from the B73 AGPv3.30 and BT×623 v3.1.1 gene sets with the R package phylostratr (Arendsee et al., 2019). Using the NCBI tree of life, we selected 6 representative genomes at each node, as well as adding recommended diverse prokaryotic taxa, for a total of 127 genomes in each analysis. We performed protein BLASTs (NCBI BLAST+) against this library of genomes with maize and again with sorghum peptides. TAI was calculated using variance-stabilized RPKM gene expression values with myTAI (Drost et al., 2018). For gene models with multiple predicted peptides, TAI was calculated with the most conserved phylostrata assigned to that locus.

We calculated maize and sorghum codon divergence phylostrata by performing reciprocal best BLAST (e-value cutoff 1E-5) for CDS from each species against the *Setaria italica* v2.2 CDS with the R package orthologr (Drost et al., 2015). Amino acids were aligned using the Needleman-Wunsch algorithm and then codon aligned with PAL2NAL before calculating substitution rates and separating into equal deciles. TDI was calculated using variance-stabilized RPKM gene expression values with myTAI (Drost et al., 2018). For gene models with multiple predicted peptides, TAI was calculated with the most conserved codon divergence strata assigned to that locus.

