## Supplemental figures for "Reconstructing the transcriptional ontogeny of maize and sorghum supports an inverse hourglass model of inflorescence development"

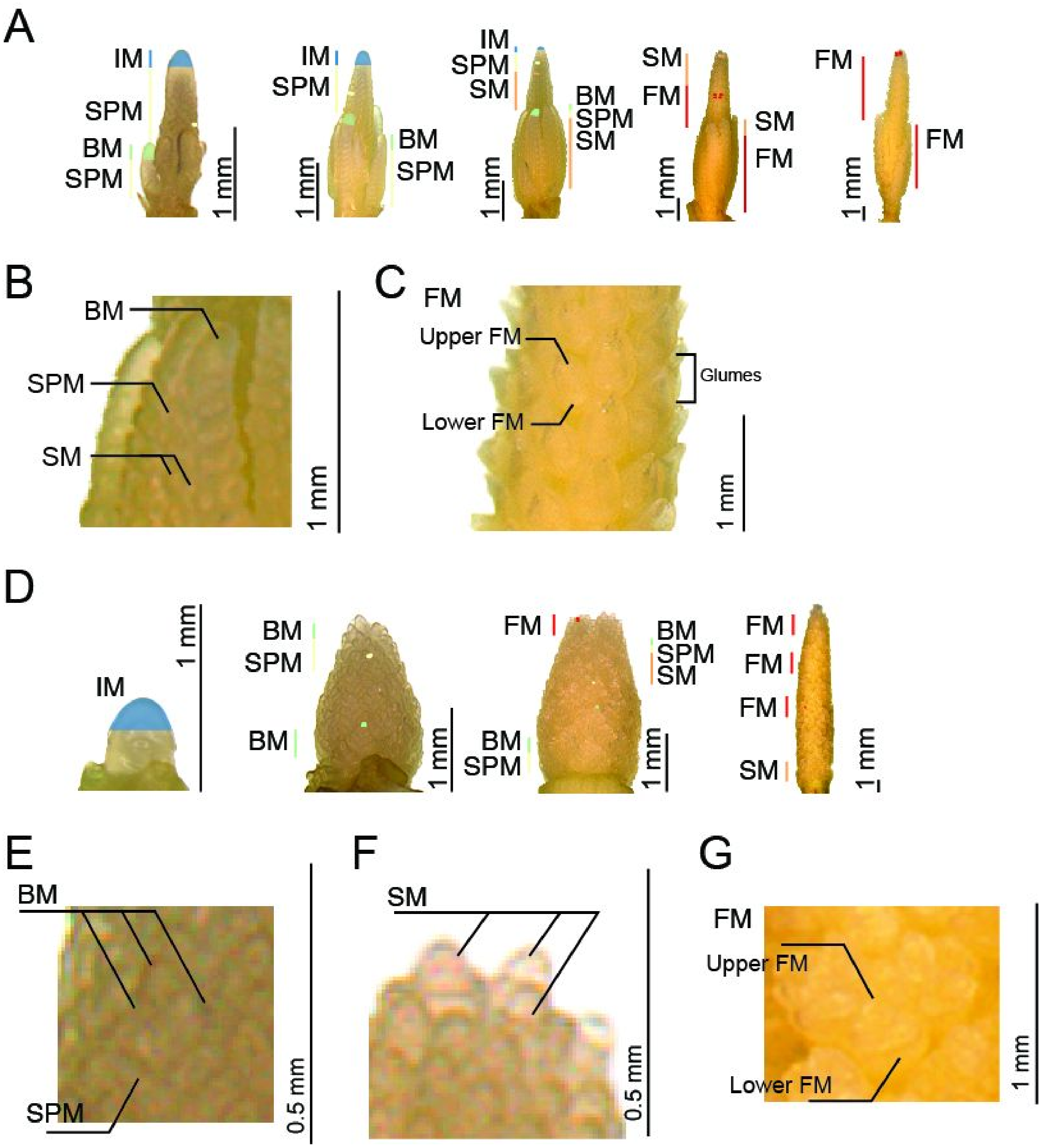


Figure 2 Supplemental figure 1. Maize and sorghum meristem identity appearances.

(A) Lateral branches pass through a sequence of identities from the IM, including BM (on branches only), SPM, SM, FM in an acropital, base-to-tip gradient in maize.
(B) Example BM, SPM, and SM on maize tassel branch.
(C) Example FMs in enclosed glumes in maize.

(D) Lateral branches pass through a sequence of identities from the IM, including BM, SPM, SM, FM in a basipetal, top-to-bottom gradient in sorghum. High order compound branching and prolonged indeterminacy are prevalent towards the base of the panicle.
(E) Example BMs and likely SPM in sorghum.
(F) Example terminal 3 SMs in sorghum.
(G) Example FMs in sorghum.

A

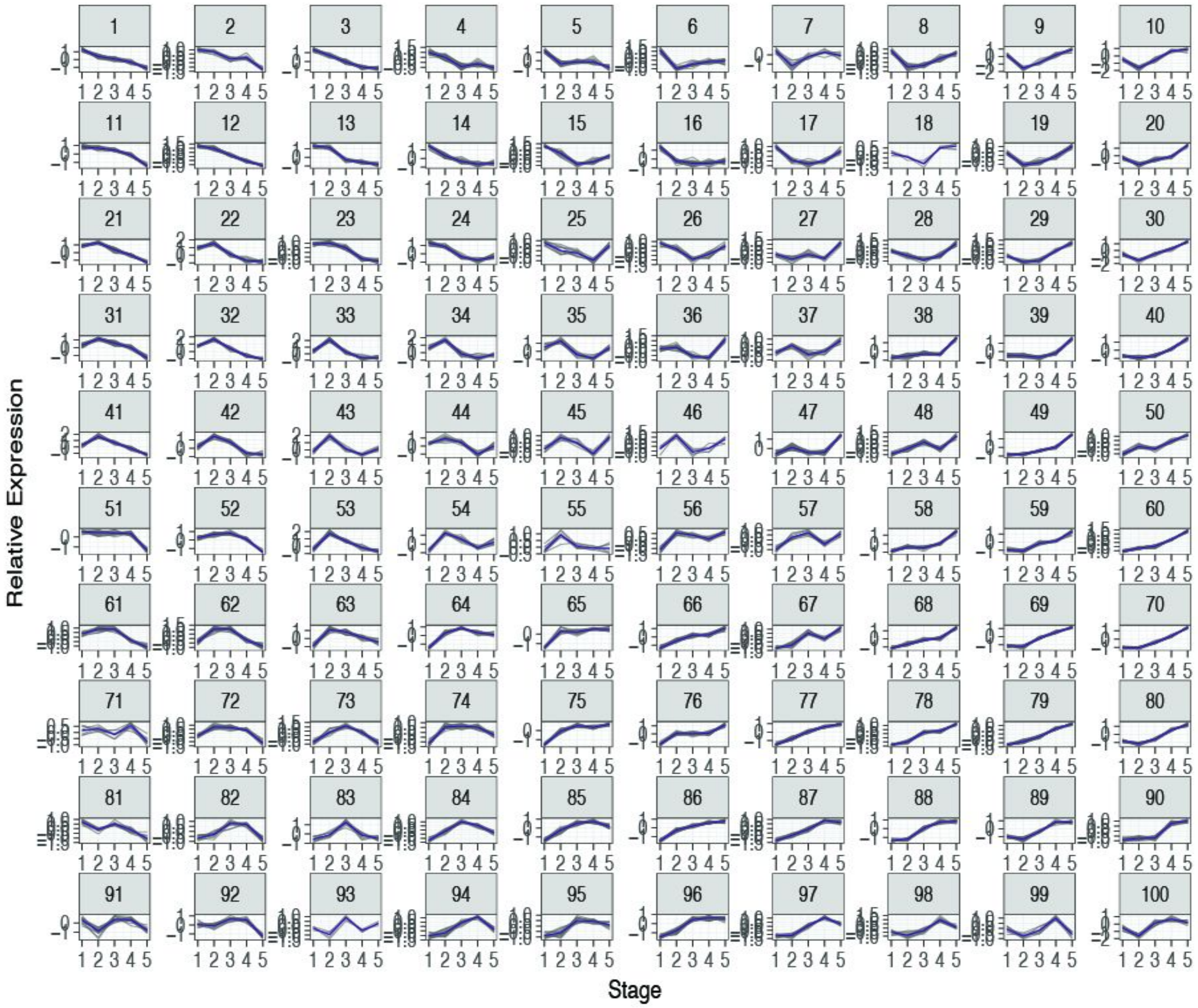


Figure 4 Supplemental figure 1. Clustered patterns of genes expressed across 5 maize tassel stages.

(A) Top 3000 most informative genes from an entrained random forest model separated into a 10 x 10 hexagonal self organizing map.

A

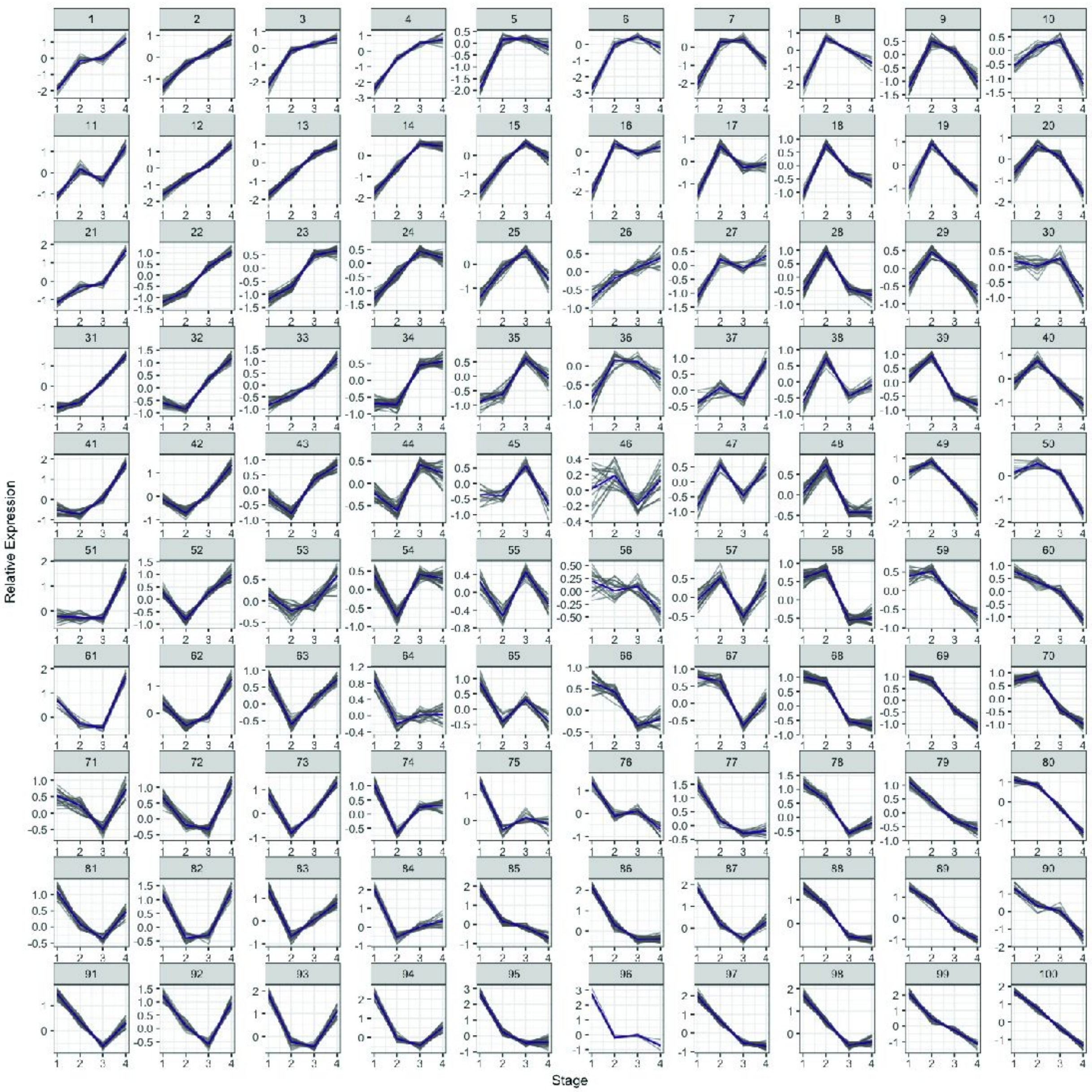


Figure 4 Supplemental figure 2. Clustered patterns of genes expressed across 4 sorghum panicle stages.

(A) Top 3000 most informative genes from an entrained random forest model separated into a 10 x 10 hexagonal self organizing map.

A

Maize Tassel Phylostrata

| Strata | Taxonomic Classification | #ZmPeptides |
|---|---|---|
| PS1 | **cellular organisms** | 21782 |
| PS2 | **Eukaryota** | 14629 |
| PS3 | **Viridiplantae** | 4180 |
| PS4 | **Streptophyta** | 3123 |
| PS5 | **Embryophyta** | 4552 |
| PS6 | **Tracheophyta** | 390 |
| PS7 | **Magnoliophyta** | 2276 |
| PS8 | **Mesangiospermae** | 1512 |
| PS9 | **Liliopsida** | 58 |
| PS10 | **Petrosaviidae** | 235 |
| PS11 | **commelinids** | 326 |
| PS12 | **Poales** | 149 |
| PS13 | **Poaceae** | 2079 |
| PS14 | **Panicoideae** | 627 |
| PS15 | **Andropogoneae** | 249 |
| PS16 | **Zea mays** | 7066 |

B

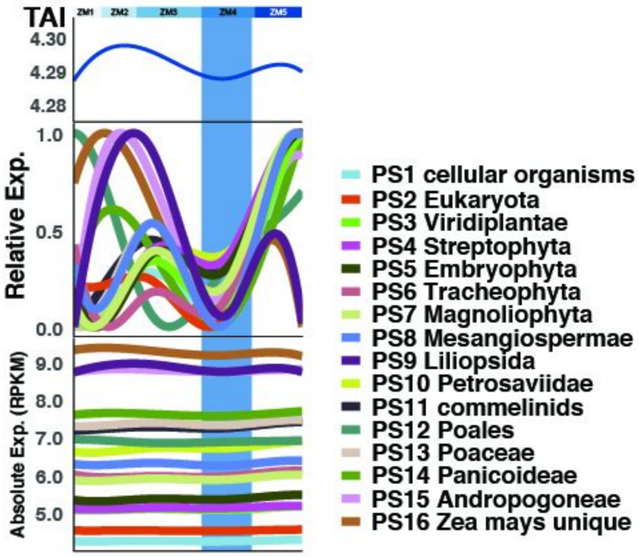


C

Sorghum Panicle Phylostrata

| Strata | Taxonomic Classification | #SbPeptides |
|---|---|---|
| PS1 | **cellular organisms** | 17038 |
| PS2 | **Eukaryota** | 3178 |
| PS3 | **Viridiplantae** | 2719 |
| PS4 | **Streptophyta** | 1961 |
| PS5 | **Embryophyta** | 3178 |
| PS6 | **Tracheophyta** | 307 |
| PS7 | **Magnoliophyta** | 1742 |
| PS8 | **Mesangiospermae** | 1411 |
| PS9 | **Liliopsida** | 32 |
| PS10 | **Petrosaviidae** | 171 |
| PS11 | **commelinids** | 252 |
| PS12 | **Poales** | 89 |
| PS13 | **Poaceae** | 2057 |
| PS14 | **Panicoideae** | 698 |
| PS15 | **Andropogoneae** | 216 |
| PS16 | **Sorghum bicolor** | 5262 |

D

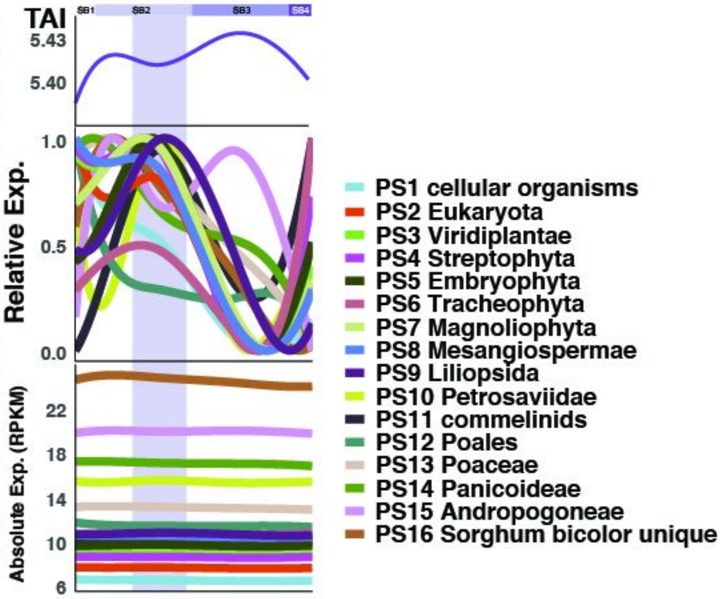


Figure 6 Supplemental figure 1. Contribution of phylostrata to TAI.

(A) Predicted maize peptides assigned to 16 phylostrata

(B) TAI (upper) is comprised by the expression values of all 16 phylostrata. Relative phylostrata contributions (middle) and absolute contributions (bottom). Hourglass-like stage, blue vertical band.

(C) Predicted sorghum peptides assigned to 16 phylostrata

(D) TAI (upper) is comprised by the expression values of all 16 phylostrata. Relative phylostrata contributions (middle) and absolute contributions (bottom). Hourglass-like stage, purple vertical band.
